# Characterization of neurite and soma organization in the brain and spinal cord with diffusion MRI

**DOI:** 10.1101/2025.02.19.638936

**Authors:** Kurt G Schilling, Marco Palombo, Atlee A. Witt, Kristin P. O’Grady, Marco Pizzolato, Bennett A Landman, Seth A. Smith

## Abstract

The central nervous system (CNS), comprised of both the brain and spinal cord, and is a complex network of white and gray matter responsible for sensory, motor, and cognitive functions. Advanced diffusion MRI (dMRI) techniques offer a promising mechanism to non-invasively characterize CNS architecture, however, most studies focus on the brain or spinal cord in isolation. Here, we implemented a clinically feasible dMRI protocol on a 3T scanner to simultaneously characterize neurite and soma microstructure of both the brain and spinal cord. The protocol enabled the use of Diffusion Tensor Imaging (DTI), Standard Model Imaging (SMI), and Soma and Neurite Density Imaging (SANDI), representing the first time SMI and SANDI have been evaluated in the cord, and in the cord and brain simultaneously. Our results demonstrate high image quality even at high diffusion weightings, reproducibility of SMI and SANDI derived metrics similar to those of DTI with few exceptions, and biologically feasible contrasts between and within white and gray matter regions. Reproducibility and contrasts were decreased in the cord compared to that of the brain, revealing challenges due to partial volume effects and image preprocessing. This study establishes a harmonized approach for brain and cord microstructural imaging, and the opportunity to study CNS pathologies and biomarkers of structural integrity across the neuroaxis.

## Introduction

The central nervous system (CNS), comprised of both the brain and spinal cord, is a highly interconnected network that facilitates a range of sensory, motor, and cognitive functions [1]. The CNS is composed of two main types of tissue, white matter (WM) and gray matter (GM). White matter is made up of a network of nerve fibers (i.e., myelinated axons) that facilitate communication between areas of the brain and spinal cord, while the gray matter is composed of neuronal cell bodies (i.e., soma) and dendrites responsible for processing and integrating neural information [2] (**Figure 1**). The brain and cord are highly interconnected, and any changes in the organization and microstructure in one region, such as the brain, could propagate and impact the structure and function of the spinal cord, or vice-versa. Therefore, it is crucial to examine both structures together when studying anatomy or pathology of the CNS. In this study, we aim to characterize the microstructural organization of both the brain and spinal cord using clinically feasible diffusion MRI (dMRI), identifying what aspects of tissue microstructure can be reliably detected and characterized in both regions.

**Figure 1.**
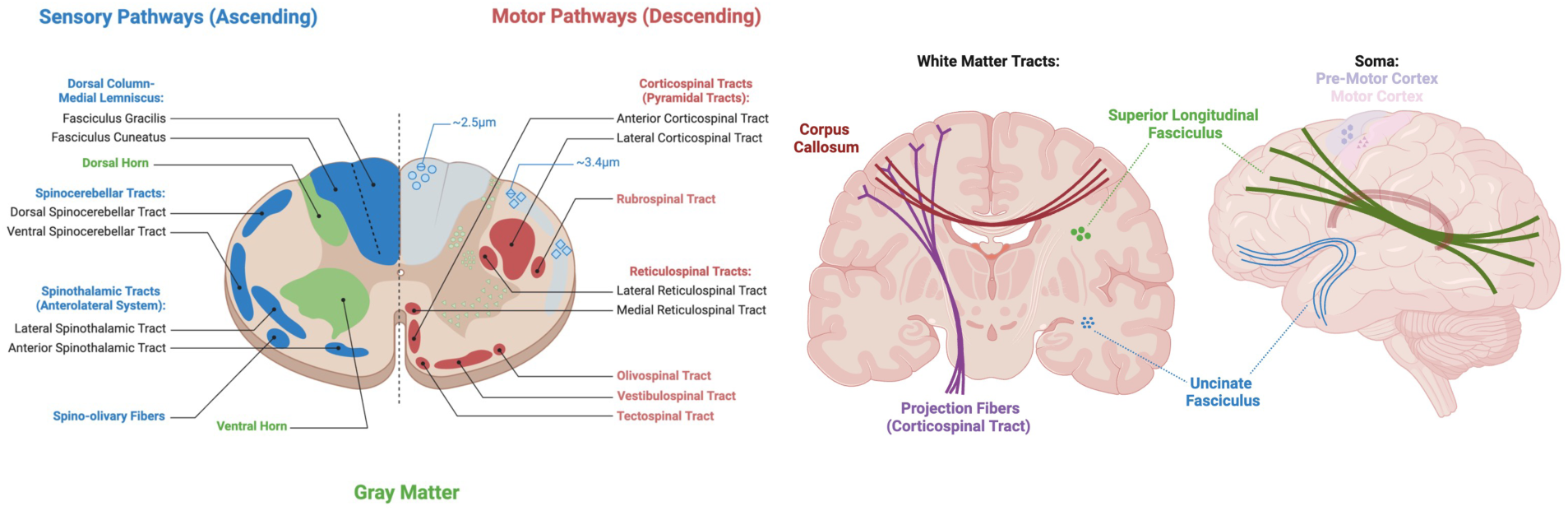
The brain and spinal cord are composed of highly organized white matter pathways, with varying neuronal densities, diameters, and orientations, as well as gray matter regions with varying cell densities and distributions. Created in BioRender. Witt, A. (2025) https://BioRender.com/s48o463

Diffusion MRI has proven to be a powerful tool to study the microstructural organization of the CNS [3, 4]. Several signal representations and microstructural models have been developed and applied to study CNS in health and disease [5, 6] (**Figure 2**).

**Figure 2.**
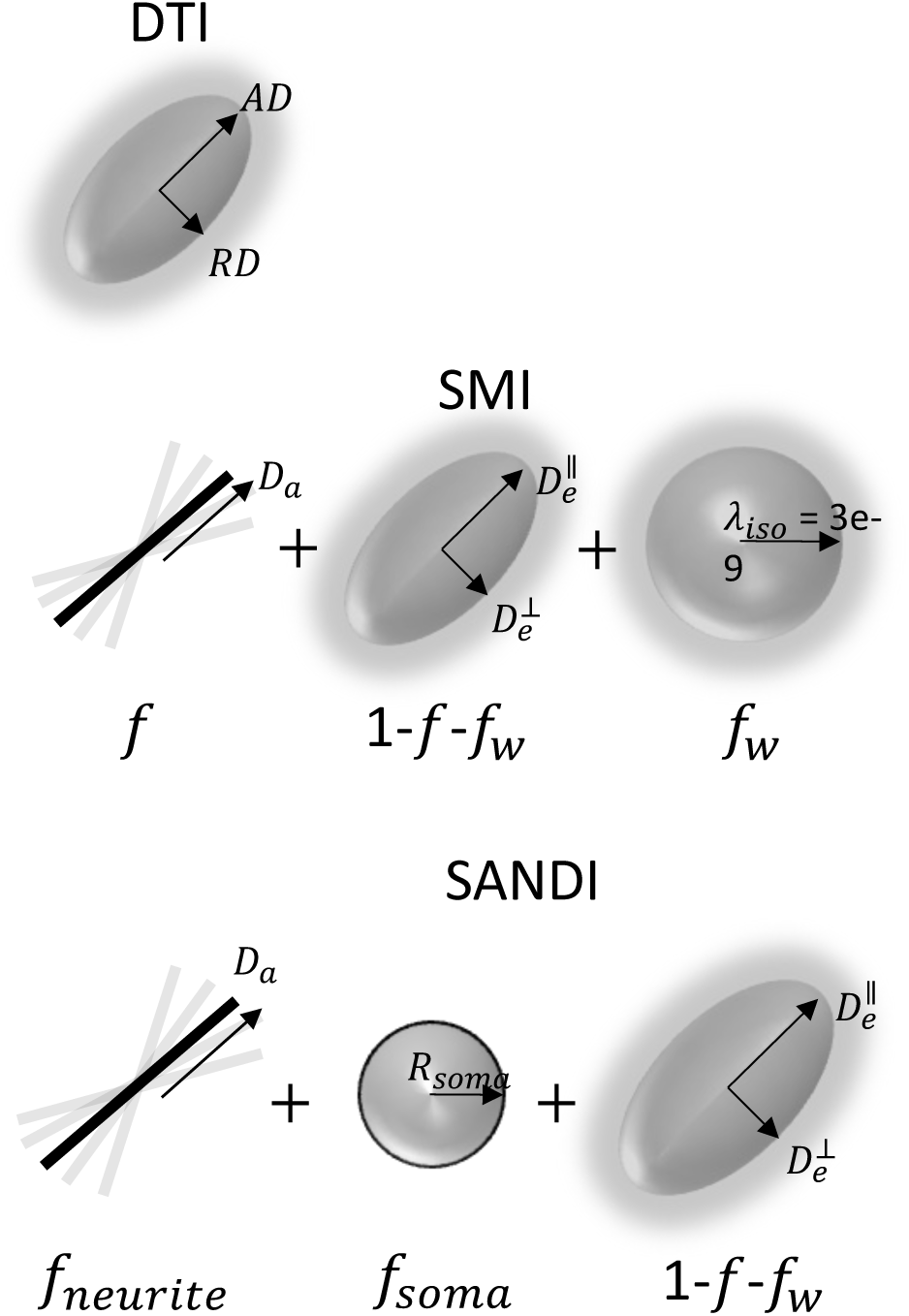
Diffusion MRI-based models of tissue microstructure. Diffusion Tensor Imaging (DTI) models the anisotropic diffusion of water in tissue as a (covariance matrix of a) 3D Gaussian distribution, from which axial (AD), radial (RD), mean diffusivities (MD), and fractional anisotropy (FA) can be derived. Standard Model Imaging (SMI) represents tisseua as zero-radius sticks embedded within an extra-axonal space described – and results in indices of axonal fraction (f), along-axon diffusivity (Da), extra-axonal parallel (De_par) and perpendicular (De_perp) diffusivities, free water (signal) fraction (fw), and a measure of fiber dispersion (p2). Soma And Neurite Density Imaging (SANDI) models water diffusion within neurites (stick-like axons and dendrites) and soma (spherical neuronal cell bodies), extending the standard model to also include soma fraction (fsoma) and soma radii (Rsoma).

Among the earliest, diffusion tensor imaging (DTI) [7, 8] captures the anisotropic diffusion of water in tissue as a (covariance matrix of a) 3D Gaussian distribution (**Figure 2, top**).

From DTI, scalar indices of diffusivity can be derived including axial (AD), radial (RD), and mean diffusivities (MD), as well as indices of orientation anisotropy like fractional anisotropy (FA) [8]. Acquisition requirements for DTI analysis are not relatively demanding, requiring only ∼15-30 diffusion-weighted images with a single b-value shell (typically b∼1000s/mm2), conditions that are readily achievable on clinical scanners [9, 10].

Because of this, DTI has been applied extensively in both the brain [11] and spinal cord [12] to study injury, neurological disease, and developmental processes in both WM and GM tissues. While DTI-based indices sensitively reflect tissue properties, such as myelination and fiber density [4], they lack specificity and are confounded by orientation and partial volume effects [9].

To overcome this limited specificity, several multi-compartmental approaches have been developed to explicitly model certain aspects of the tissue environment [5, 6]. Among others [13–18], models such as neurite orientation dispersion and density imaging (NODDI) [19], multi-compartment spherical mean technique (SMT) [20, 21], or White Matter Tract Imaging (WMTI) [22, 23], have become popular in both the brain and spinal cord because they can be employed using a clinically feasible two-shell (two b-values) acquisition. While there are unique differences in constraints, assumptions, and derived indices, many of these models fall under the umbrella of the so-called *Standard Model* of neuronal tissue [5]. In Standard Model Imaging (SMI) (**Figure 2, middle**), axons are represented as impermeable zero-radius sticks arranged in coherent bundles and embedded within an extra-axonal space described by an axially symmetric diffusion tensor, along with a third cerebral spinal fluid (or free water) compartment. Thus, SMI results in indices of axonal (or signal) fraction (f), along-axon diffusivity (Da), extra-axonal parallel (De_par) and perpendicular (De_perp) diffusivities, free water (signal) fraction (fw), and a measure of fiber dispersion (p2). These indices have been shown to be highly specific to normal anatomical variation [24] and disease processes such as demyelination (De_perp) [25, 26], axonal loss (f) [25], and axonal beading (Da) [27], with reproducible and robust estimates on clinical scanners [28]. However, SMI has not yet been demonstrated in the spinal cord, where descriptions of normal variation and reproducibility could well-compliment the same measures derived in the brain. Additionally, the standard model is based on geometrical assumptions specific to white matter and may not be entirely appropriate for gray matter tissue.

More recently, the Soma and Neurite Density Imaging (SANDI) model [29] has emerged as a promising technique for studying both white matter and gray matter architecture in the CNS. By distinguishing between water diffusion within neurites (stick-like axons and dendrites) and soma (spherical neuronal cell bodies), SANDI provides a more comprehensive characterization of tissue microstructure than previously mentioned models, extending the standard model to also include soma fraction (fsoma) and soma radii (Rsoma) estimates (**Figure 2, bottom**). However, this comes at the cost of a significantly increased scan time, requiring a minimum of 5 diffusion shells, with b-values 6-10x higher than those typical of clinical acquisitions. In the original paper [29], the SANDI model was fit on data with b-values up to 10,000s/mm2 acquired on a high-performance scanner, showing white and gray matter contrasts that paralleled those obtained from myelo-architectural and cyto-architectural stains. More recently, Genc et al., [30] revealed that these maps were highly repeatable and reproducible using much lower b-values of b=6000s/mm2, but still acquired using high performance systems. Finally, Schiavi et al. [31] demonstrated the feasibility of acquiring SANDI metrics on a clinical scanner, highlighting biases due to noise and acquisition schemes, but resulting in reliable and reproducible measures in both white and gray matter. However, again, these were demonstrated in the brain, and feasibility of SANDI modeling has not been done on the cord.

Motivated by these works in the brain, and the desire to study both the brain and spinal cord in parallel with a harmonized acquisition, we aim to characterize neurite and soma organization in the brain and spinal cord using diffusion MRI on a clinical scanner. In agreement with existing literature, we show that DTI, SMI, and SANDI are feasible, reproducible, and result in biologically meaningful measures of the brain, but address challenges in image acquisition and image processing [32, 33] to demonstrate similar application of these models in the cord. We first describe the acquisition which enables application of these models in the gray and white matter, then show and assess data quality and resulting indices in both the brain and cord. Next, we assess reliability and repeatability of these models, and finally compare model indices to known anatomy and expected contrast within and between white and gray matter tissues.

## Methods

### Brain and Cord - acquisition

Eleven healthy controls participated in this study, with six scanned twice for scan-rescan reproducibility assessment. All experiments were performed on a 3.0T whole body MR scanner (Philips dStream Ingenia, Best, Netherlands). A two-channel body coil was used for excitation and a 16-channel SENSE neurovascular coil was used for reception.

The maximum gradient strength of the system was 80 mT/m at a slew rate of 100 mT/m/s.

All data were acquired under a protocol approved by the local institutional review board (IRB #111087) and informed consent was obtained prior to the study.

Brain imaging consisted of a T1-weighted image using a three-dimensional (3D) T1-MPRAGE image (TR=6.3ms, inversion time = 1060ms, TE = 2.9ms, flip angle 8 degrees, spatial resolution 1×1×1mm3, acquisition time = 5m37s). The diffusion protocol consisted of a pulsed-gradient spin-echo sequence with single-shot echo planar imaging (EPI) readout with the following parameters: repetition time 4400 ms, echo time 100 ms, pulse duration 24 ms, pulse separation 60 ms, spatial resolution 2.5 x 2.5 x 2.5mm3, 62 slices, axial orientation, SENSE=2.2, Multiband=2, partial Fourier =0.70, with 7 (non-zero) b-values of 0/100/500/1000/2000/3000/4000/6000 with 22/8/8/16/24/32/40/48 measurements per shell, acquired with phase encoding in the anterior-posterior direction, with a scan of 6 b=0s/mm2 images with reversed phase encoding. The total scan time for brain diffusion data was 16.5 minutes. We note that the sequence was separated into three sequential acquisitions, with constant pre-scan settings, because the software did not allow >128 diffusion-weighted volumes in a single acquisition. Scans were simply concatenated prior to preprocessing.

Cord imaging consisted of a high-resolution (0.65 × 0.65 × 5 mm^3^) multi-slice, multi-echo gradient echo (mFFE) anatomical image [34] (TR/TE/ΔTE = 753/7.1/8.8 ms, α = 28°, number of slices = 14, 6:12 min) for co-registration and to serve as a reference image for segmentation. The diffusion protocol was matched to the brain, and consisted of a pulsed-gradient spin-echo sequence with single-shot EPI readout with the following parameters: repetition time 4400 ms, echo time 100 ms, pulse duration 24 ms, pulse separation 51 ms, spatial resolution 1.1 x 1.1 x 5mm3, 18 slices, axial orientation, SENSE=1.8, Multiband=None, partial Fourier = 0.69, with 7 (non-zero) b-values of 0/100/500/1000/2000/3000/4000/6000 with 22/8/8/16/24/32/40/48 gradient directions performed per shell, acquired with phase encoding in the left-right direction, with a scan of 6 b=0s/mm2 images with reversed phase encoding. All images were centered at the C3/C4 intervertebral disc. Reduced field-of-view was applied using an outer volume suppression technique [35] and fat suppression was achieved using SPIR. The acquisition was not cardiac triggered in order maximize acquisition per time [36]. The total scan time for cord diffusion data was 16.5 minutes. Again, the sequence was separated into three sequential acquisitions, with constant pre-scan settings, and concatenated prior to preprocessing.

### Brain and Cord - Preprocessing

Brain data preprocessing started with running FreeSurfer 6.0 [37, 38] in order to derive 24 regions-of-interest (ROIs) from the lobe-based parcellation including 12 gray matter labels (bilateral Frontal, Parietal, Occipital, Temporal, Cingulate, Subcortical) and the corresponding 12 white matter labels. The diffusion dataset was preprocessed using a combination of FSL and MRTrix3 software, and included MPPCA denoising [39], Gibbs-Ringing correction [40], and correction of motion, eddy currents, and susceptibility distortion[41, 42], followed by a final round of denoising using Patch2Self[43], which can be applied at any point in the preprocessing pipeline. Noise maps were estimated from the MPPCA denoising process.

Spinal cord data preprocessing utilized tools from the Spinal Cord Toolbox [44]. Preprocessing started with vertebral labelling on the structural mFFE and subsequent registration to the PAM50 template, resulting in 24 ROIs, including 6 gray matter labels (bilateral Ventral Horns, Intermediate Zones, Ventral Horns) and 6 white matter labels (bilateral Dorsal Columns, Lateral Columns, Ventral Columns), for each cervical level 3 and cervical level 4. Diffusion preprocessing included MPPCA denoising [45], Gibbs-Ringing correction [40], motion correction (using SCT *sct_dmri_moco* [46] including slice-wise regularized registration along the SI-axis, iterative averaging of the target volumes, and group-wised alignment of DWIs with grouping g=8), followed by Patch2Self denoising [43]. Noise maps were estimated from the MPPCA denoising process. All data was quality checked at every point in the process, with manual inputs to vertebral alignment, masking, and alignment parameters if needed.

### Microstructure Modeling

The DTI, SMI, and SANDI models were fit to the preprocessed data for both brain and cord.

DTI was fit using FSL toolbox command *dtifit*, using only b-values b<=2000 (b=100, 500, 1000, 2000) and simultaneously fitting both the Diffusion and Kurtosis tensors [47] using linear least squares on the log-transformed signal. This resulted in maps of FA, MD, AD, and RD, describing the degree of diffusion anisotropy, the mean of the three eigenvalues of the diffusion tensor, the diffusion parallel to the principal diffusion direction, and the diffusion perpendicular to the principal diffusion direction, respectively. Finally, from the kurtosis tensor [47], the mean kurtosis (MK) describing the average of the diffusion kurtosis along all directions. Kurtosis describes non-Gaussianity of the diffusion process and has been interpreted as a more complex diffusion pattern within the imaging voxel (due to hindrances and restrictions).

SMI fitting was performed using the Standard Model Imaging Matlab toolbox (https://github.com/NYU-DiffusionMRI/SMI) [28] providing b-values b<=3000 (b=100, 500, 1000, 2000, 3000) and the estimated noise map for Rician bias correction prior to fitting. This toolbox used root-mean-squared error (RMSE) machine-learning based estimators for parameter estimation [48]. This resulted in maps characterizing intracellular space including axonal fraction (f) describing the relative signal contribution of intra-axonal space, intra-axonal diffusivity (Da) along axons, and orientational coherence parameter (p2). Maps characterizing extracellular space included the radial diffusivity (De_perp) and axial diffusivity (De_par). And finally, the free water fraction (fw) describing the signal fraction of an isotropic free water component.

To fit the SANDI model we used the SANDI Matlab Toolbox (https://github.com/palombom/SANDI-Matlab-Toolbox-Latest-Release), using as input the respective preprocessed diffusion data and noise maps for the brain and cord. The toolbox similarly uses machine-learning based estimators for parameter estimation. We fixed the intra-soma diffusivity to 3 um2/ms, and upper bounds for Din and De to 3 um2/ms, and soma radius upper bound automatically set by default to a maximum value given the diffusion time and intra-soma diffusivity.

### Analysis

To evaluate image quality, the apparent signal-to-noise ratio (SNR) (the SNR after the adopted image preprocessing) was calculated following [31], using two consecutive b=0 images:

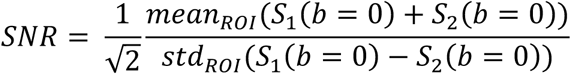

where S1 and S2 are the first two consecutive b=0 images. The SNR was calculated in three regions-of-interest (ROIs), including white matter, gray matter, and their sum, for both the brain and spinal cord.

The reliability of each diffusion metric, for all models, in both the brain and cord, was evaluated using the three commonly used statistical measures in medical imaging [31, 49–51]: Pearson correlation r, the test-retest variability (TRV), and intraclass correlation coefficient (ICC). The Pearson correlation r was computed using the MatLab *corrcoeff* function using the test-retest values of the 24 brain ROIs (12 WM and 12 GM) and 24 cord ROIs (12 WM and 12 GM). As in [52], we interpreted r less than 0.4 as weak correlation, r between 0.4 and 0.69 as moderate correlation, r between 0.70 and 0.89 as strong correlation, and r greater than 0.90 as very strong correlation.

TRV was compute across the 24 ROIs (for both brain and cord) for each parameter θ as:

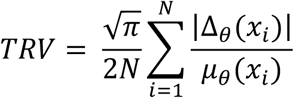

where Δ_θ_(x_i_) and μ_θ_(x_i_) were the difference and average of the test and retest estimates of the parameter θin the *i*th ROI x_i_.

Finally, ICC was calculated using a two-way mixed effects, single measurement model, using the Matlab ICC function available through Mathworks File Exchange [53]. As in [49, 50], we interpreted ICC less than 0.5 as poor reliability, ICC between 0.5 and 0.75 as moderate reliability, ICC between 0.75 and 0.9 as good reliability, and ICC greater than 0.9 as excellent reliability.

Finally, to assess variation across the brain and cord, we plot the distributions of parameters, for each model, across the 12 WM and 12 GM regions of interest.

## Results

### Acquisition is feasible, resulting in high image quality, even at high b-values

Figure 3 shows example diffusion weighted images in the brain and cord for 5 randomly selected subjects, at all b-values. In the brain, contrast is observed across white matter regions with varying orientations, with high signal in pathways orthogonal to the diffusion sensitization directions, even at b=6000. In the cord, contrast is observed across white and gray matter tissue, with the gray matter ‘butterfly’ visible at lower diffusion-weighted signal indicative of less restrictions and greater diffusivity. However, there is significant partial volume between the tissues due the small size of the intra-cord structures. Regardless, signal remains when diffusion sensitization is orthogonal to the cord, even at b=6000. **Table 1** reports the apparent SNR in the cord for all subjects and rescans. In line with existing literature in the brain, our apparent SNR in WM and GM of 42.8 and 42.8, respectively. The SNR of the cord was lower, with an average value in WM and GM of 16.5 and 24.0, respectively, in line with the lower voxel sizes acquired in the cord.

**Figure 3.**
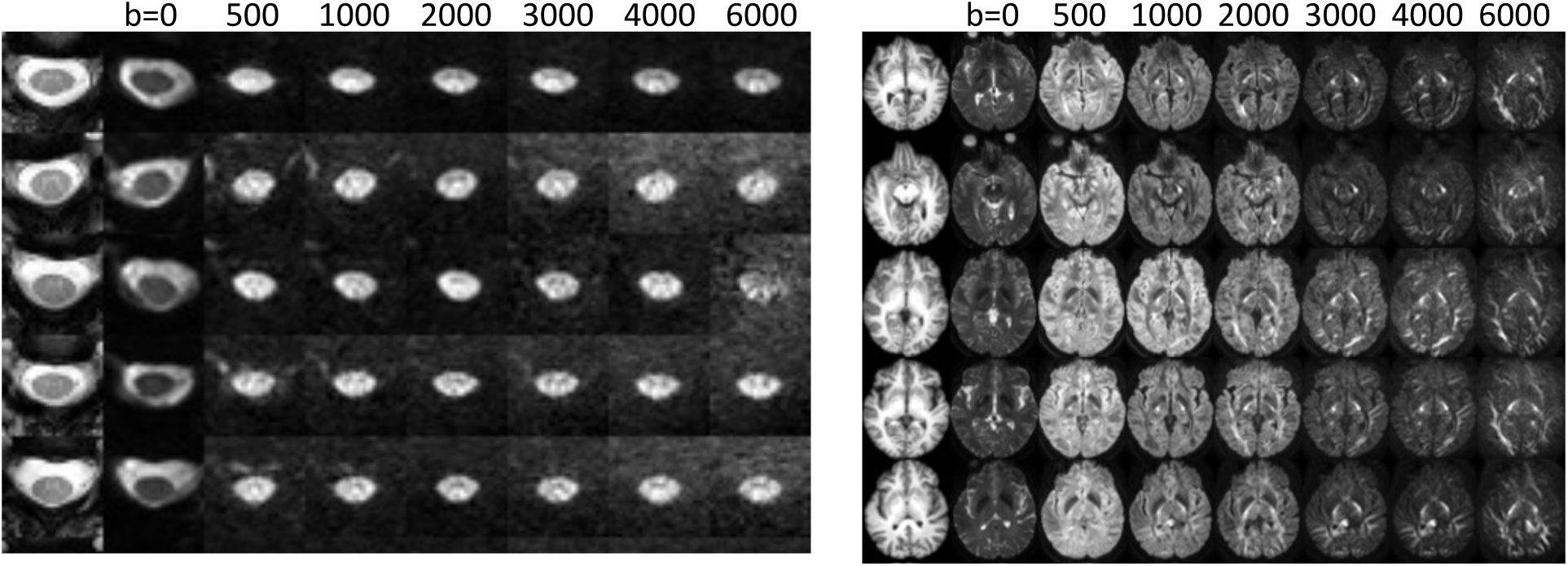
Data acquisition is feasible on a clinical scanner in the brain and spinal cord. A representative axial slice of 5 subjects is shown for the brain and cord, along with a diffusion weighted image sensitized along the Left/Right direction at b-values ranging from 0 to 6000. Contrast is visible within and across tissue types.

**Table 1.**
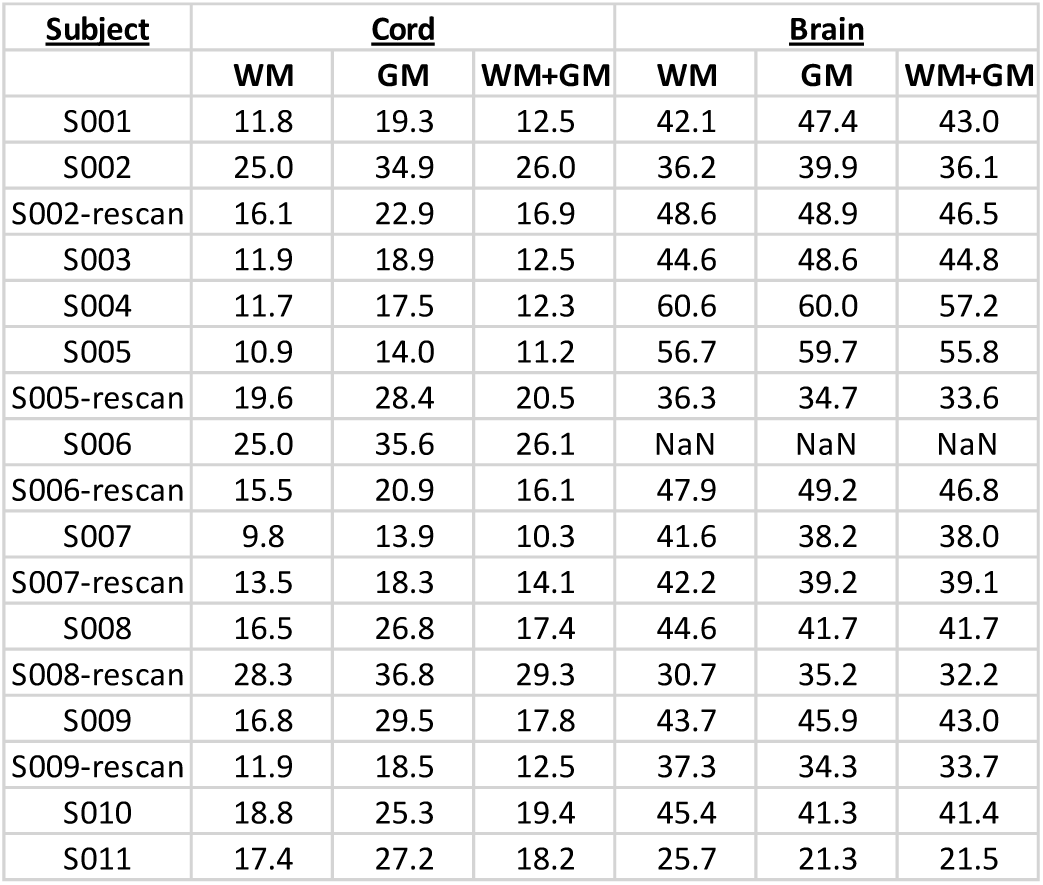
Apparent SNR for WM and GM in the brain and cord.

| <u>Subject</u> | <u>Cord</u> |  |  | <u>Brain</u> |  |  |
| --- | --- | --- | --- | --- | --- | --- |
|  | WM | GM | WM+GM | WM | GM | WM+GM |
| S001 | 11.8 | 19.3 | 12.5 | 42.1 | 47.4 | 43.0 |
| S002 | 25.0 | 34.9 | 26.0 | 36.2 | 39.9 | 36.1 |
| S002-rescan | 16.1 | 22.9 | 16.9 | 48.6 | 48.9 | 46.5 |
| S003 | 11.9 | 18.9 | 12.5 | 44.6 | 48.6 | 44.8 |
| S004 | 11.7 | 17.5 | 12.3 | 60.6 | 60.0 | 57.2 |
| S005 | 10.9 | 14.0 | 11.2 | 56.7 | 59.7 | 55.8 |
| S005-rescan | 19.6 | 28.4 | 20.5 | 36.3 | 34.7 | 33.6 |
| S006 | 25.0 | 35.6 | 26.1 | NaN | NaN | NaN |
| S006-rescan | 15.5 | 20.9 | 16.1 | 47.9 | 49.2 | 46.8 |
| S007 | 9.8 | 13.9 | 10.3 | 41.6 | 38.2 | 38.0 |
| S007-rescan | 13.5 | 18.3 | 14.1 | 42.2 | 39.2 | 39.1 |
| S008 | 16.5 | 26.8 | 17.4 | 44.6 | 41.7 | 41.7 |
| S008-rescan | 28.3 | 36.8 | 29.3 | 30.7 | 35.2 | 32.2 |
| S009 | 16.8 | 29.5 | 17.8 | 43.7 | 45.9 | 43.0 |
| S009-rescan | 11.9 | 18.5 | 12.5 | 37.3 | 34.3 | 33.7 |
| S010 | 18.8 | 25.3 | 19.4 | 45.4 | 41.3 | 41.4 |
| S011 | 17.4 | 27.2 | 18.2 | 25.7 | 21.3 | 21.5 |

### DTI and multi-compartment models show reasonable contrast between and within tissues

Figure 4 shows DTI parameter maps for 5 subjects. In the brain, white and gray matter contrast is visible in all maps, with values typical of that in the literature [8]. In the cord, white and gray matter contrast is most visible in FA and AD maps, although partial volume effects are apparent. White matter cord values of FA, MD are qualitatively similar to that of the brain, although AD appears to be greater in the cord.

**Figure 4.**
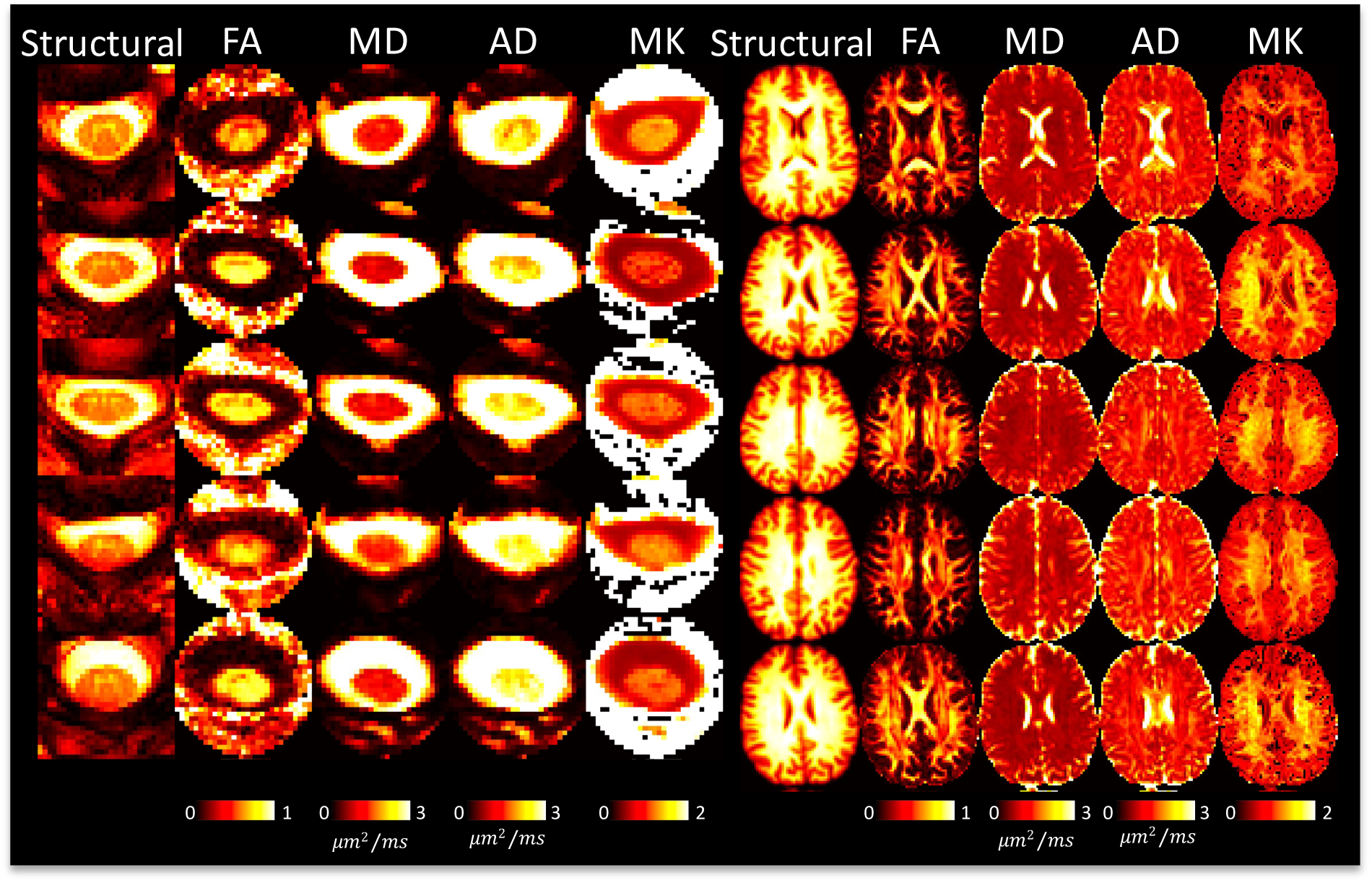
DTI shows reasonable contrast between and within tissues in the brain and spinal cord. A representative axial slice of 5 subjects is shown for the brain and cord, showing the structural image and the corresponding DTI-based contrasts.

Figure 5 shows SMI parameter maps for 5 subjects. Again, in agreement with the literature, not only is there white matter/gray matter contrast, but variation across white matter, particularly within neurite fraction (f) and orientation dispersion (p2). Freewater fraction (fw) is 0 or near-0 in white matter. Figure 5 displays the first SMI-derived maps of the spinal cord. Here, some contrast is noticeable between tissues, with gray matter visually having reduced neurite fraction (f) and De_perp, although the full gray matter butterfly is not clearly delineated. The freewater fraction is non-negligible throughout the cord, particularly on voxels neighboring the CSF, but also throughout the entire tissue, further suggesting partial volume effects in the cord.

**Figure 5.**
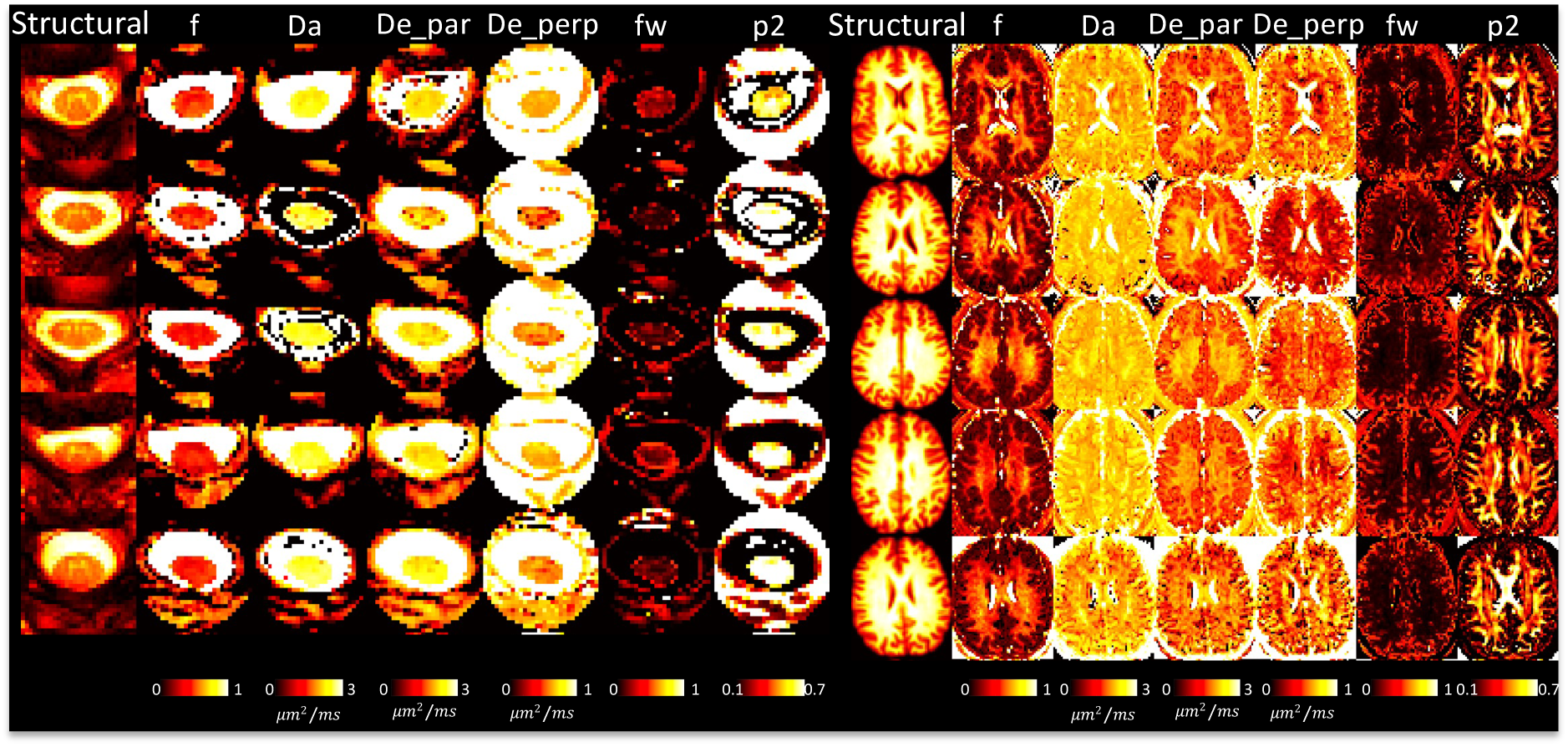
SMI shows reasonable contrast between and within tissues in the brain and spinal cord. A representative axial slice of 5 subjects is shown for the brain and cord, showing the structural image and the corresponding SMI-based contrasts.

Figure 6 shows SANDI model parameters for 5 subjects. Neurite fraction and diffusivities in the white matter of the brain well-match those of the simpler SMI model. Unique from SMI, we now have soma fraction and soma radii, both of which are increased in gray matter, with some variation across lobes. SANDI in the cord suggests less visible contrast between tissue types, particularly with soma size and fraction, although the cord-averaged value appears to be in line with that of the brain white matter. Some intra-cord contrast is visible with De and fneurite parameters, although again, partial volume effects are clear, and the GM shape is not clearly delineated. All descriptors of neurites and soma are relatively homogenous across the cord, and qualitatively similar to the values in WM of the brain.

**Figure 6.**
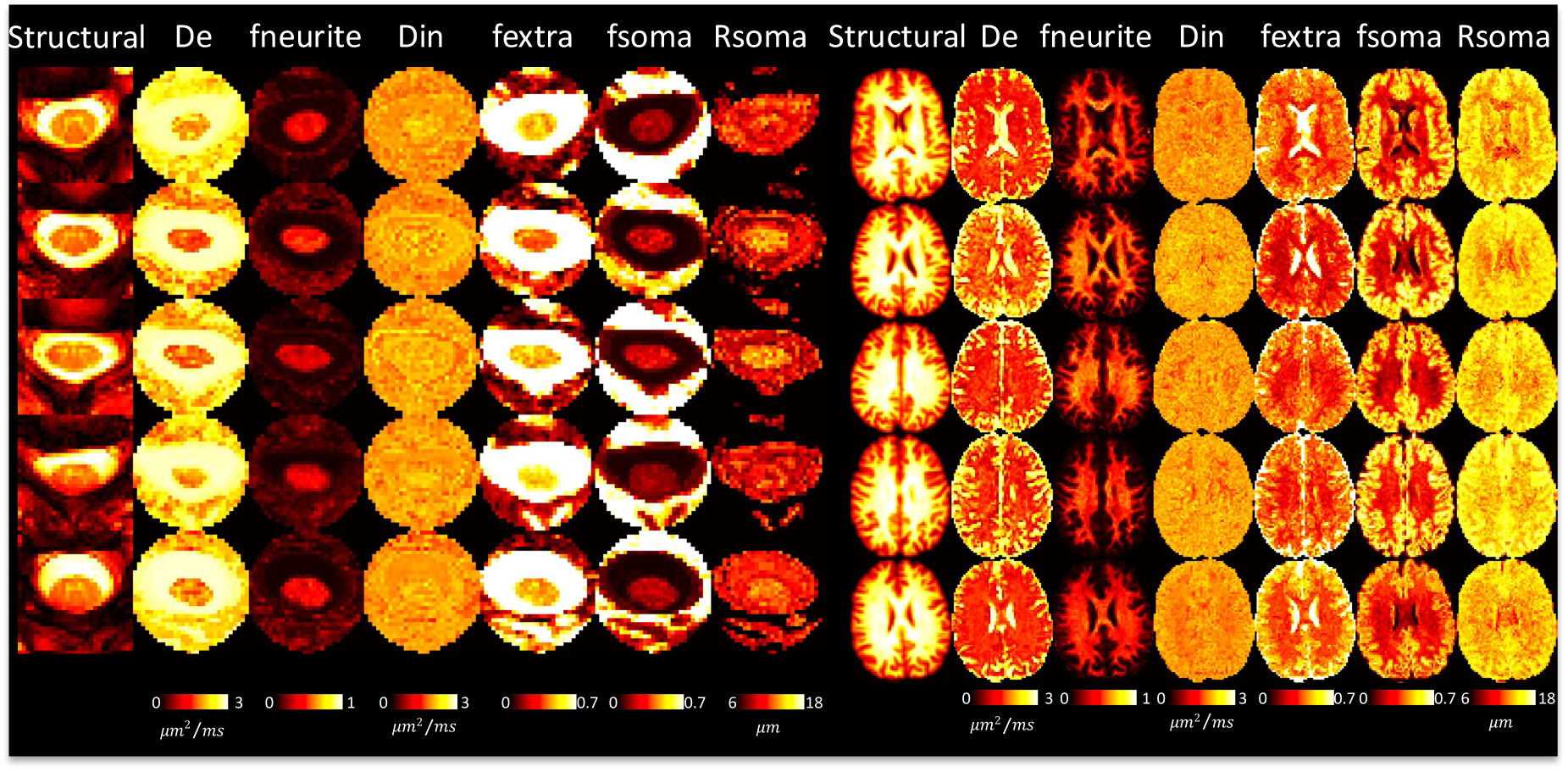
SANDI shows reasonable contrast between and within tissues in the brain and spinal cord. A representative axial slice of 5 subjects is shown for the brain and cord, showing the structural image and the corresponding SMI-based contrasts.

### DTI is highly reproducible in the brain, SMI and SANDI reproducibility is comparable to that of DTI

To assess reproducibility of the microstructural measures in the brain provided by SMI and SANDI, we compared test-retest reproducibility with those obtained from DTI (the gold standard in clinical practice) as done in [49]. This is shown as scatter plots and Pearson correlation coefficient r shown in Figure 7. Pearson r measures range from 0.63-0.98 for DTI-derived measures, with the highest for FA, describing orientation coherence. Both SMI and SANDI have similar reproducibility, with Pearson r ranging from 0.54-0.97 and 0.45-0.98, respectively. For both, neurite fractions are highly reproducible, as are orientation dispersion (p2) from SMI and soma fraction (fsoma) from SANDI, with the lowest reproducibility associated with extracellular diffusivities.

**Figure 7.**
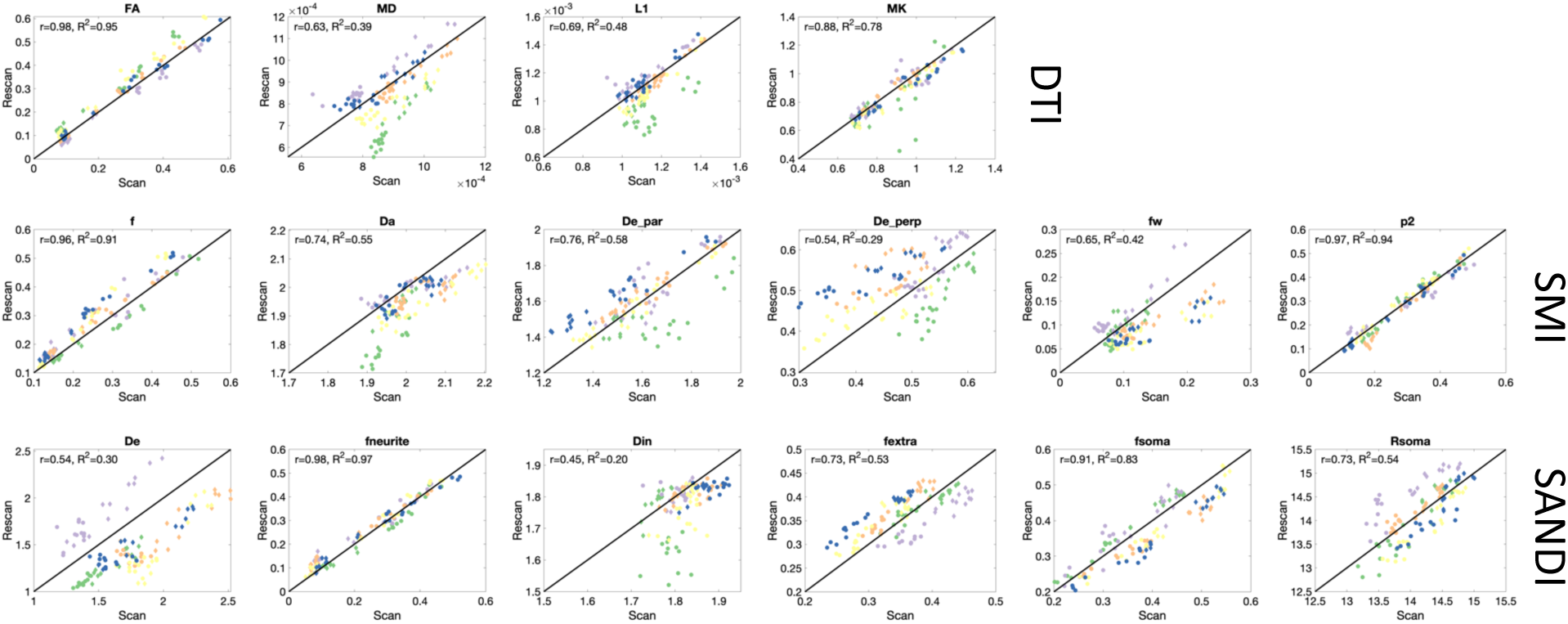
Repeatability of microstructural measures in the brain. Scatter plots of scan-rescan microstructural measures extract from 12GM and 12WM regions are shown for DTI, SMI, and SANDI models. Points are colored based on subjects.

### DTI is moderately reproducible in the cord, SMI and SANDI reproducibility less than that of DTI

Similarly, reproducibility of diffusion measures in the cord are assessed using test-retest scans and are shown in scatter plots in Figure 8. Overall, reproducibility is reduced but remains moderate for most measures. DTI ranges from 0.33-0.75 with lowest reproducibility of MD, SMI ranges from 0.3-0.83 with lowest also for extracellular diffusivity measures, and SANDI from 0.12-0.64, with low reproducibility for intracellular diffusivity (Din) and fsoma (fs).

**Figure 8.**
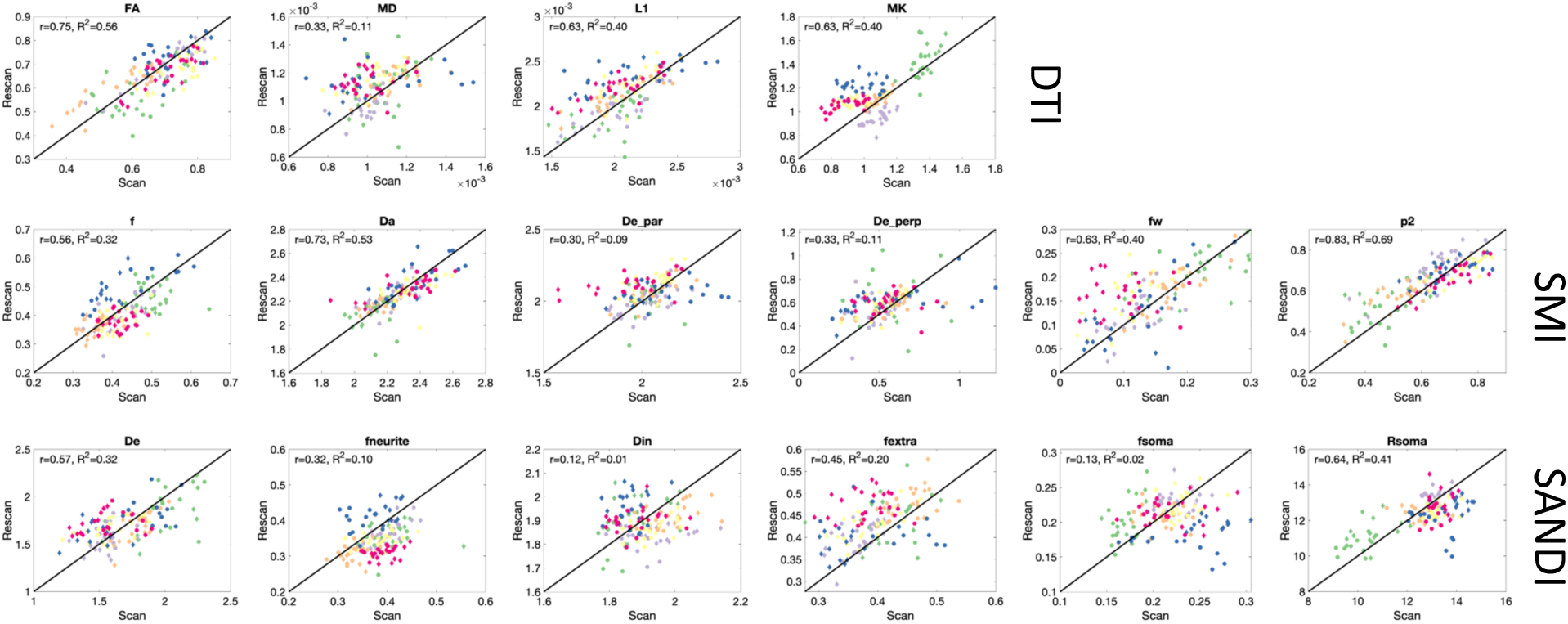
Repeatability of microstructural measures in the cord. Scatter plots of scan-rescan microstructural measures extract from 12GM and 12WM regions are shown for DTI, SMI, and SANDI models. Points are colored based on subjects.

### Measures show low test-retest variability, moderate ICC

TRV and ICC for DTI, SMI, and SANDI models, in the brain and cord, are shown in **Table 2**. Test-retest variability of SANDI and SMI is on par with that of DTI. TRV ranges of DTI, SMI, and SANDI in the cord are 7.5-11.3, 3.11-29.9, and 3.9-12.0 while those in the brain are 5.2-13.1, 3.0-28.6, and 2.0-19.69. Thus, while Pearson r across regions is low, the metrics themselves show little deviation between scan and rescan, and similar values in the spinal cord as in the brain. In general, ICC decreases from DTI, to SMI, to SANDI, and decreases from brain to cord (although this is not true across all measures), with particularly low reliability (low ICC) of Din and fsoma in the cord.

**Table 2.**
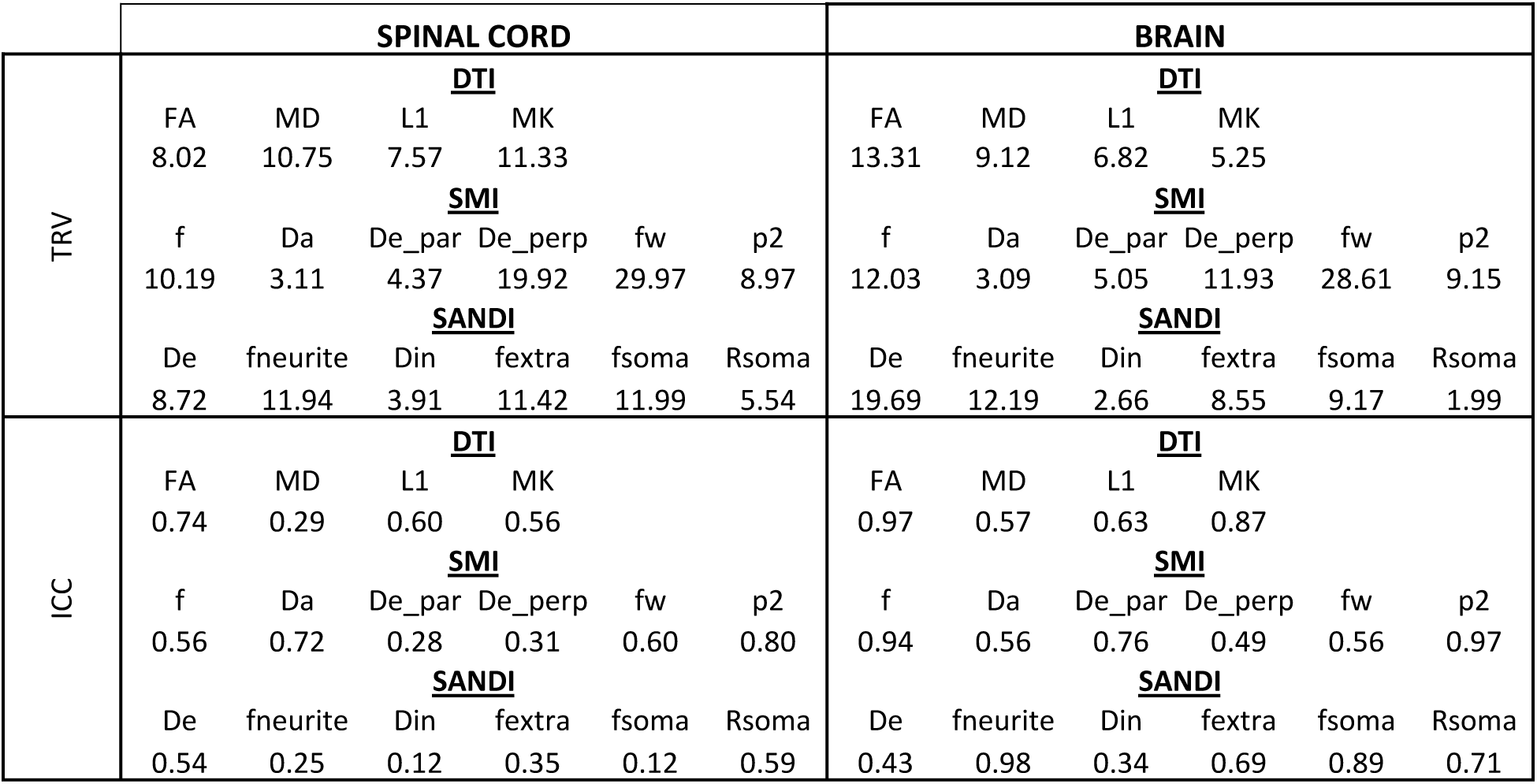
Reproducibility and Intraclass Correlation Coefficient of microstructural measures in the brain and cord.

### Soma and Neurite Density and organization show contrast within and between tissue types in the brain

Figure 9 shows distributions across the sample population of microstructural measures across the brain, for both WM (Figure 9, left) and GM (Figure 9, right) regions. For all models, several measures show clear trends across white and gray matter regions. For example, DTI measures of FA and L1, SMI measures of f, De_par, and p2, and SANDI measures of fneurite, fextra, and fsoma show clear variation across white matter. These same metrics additionally show trends across gray matter regions, with SANDI offering additional soma-based contrast that varies across the cortex. In all cases, measures are more similar to contralateral regions than across regions of the same hemisphere.

**Figure 9.**
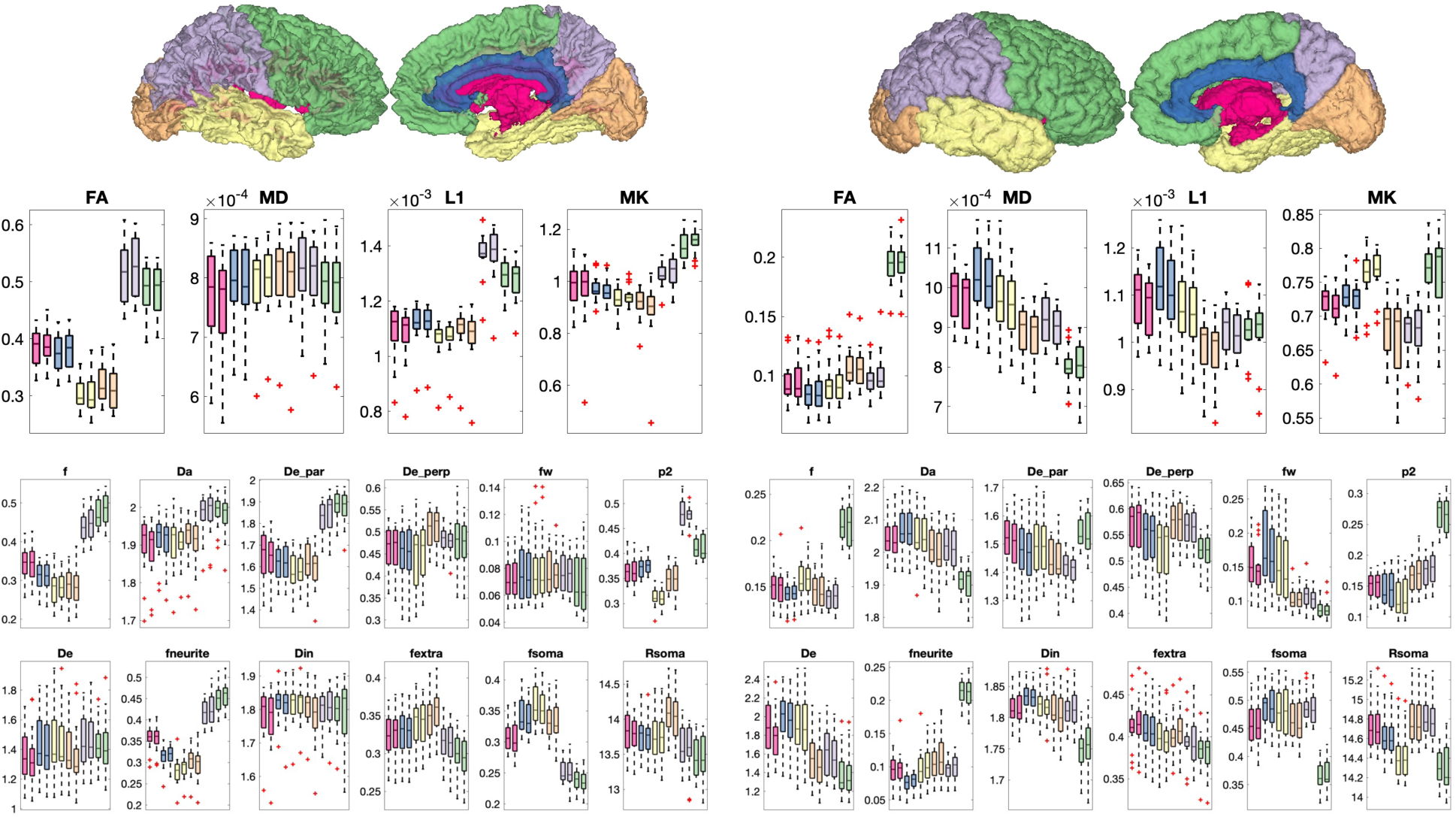
DTI (top), SMI (middle), and SANDI (bottom) show contrast between and within white matter regions (left) and gray matter regions (right). Plots are colored based on location in the lobes of the brain and corresponding white matter regions.

### Soma and Neurite Density and organization show contrast within tissue types in the cord

Figure 10 shows regional values of DTI, SMT, and SANDI-derived microstructural measures across the cord, for both WM (Figure 10, left), and GM (Figure 10, Right). As in the brain, all models show regional heterogeneity in both tissue types, although with less visible contrast than the brain. For example, DTI (L1, MD), SMI (Da, f, De_perp, fw), and SANDI (Din, fextra) show contrast across white matter pathways, while most measures show contrast across gray matter regions. All SMI measures show clear gray matter trends, and the soma-based SANDI measures also show differences **across** the ventral horn, intermediate zone, and dorsal horns. Again, measures are more similar to contralateral regions than across regions of the same hemisphere.

**Figure 10.**
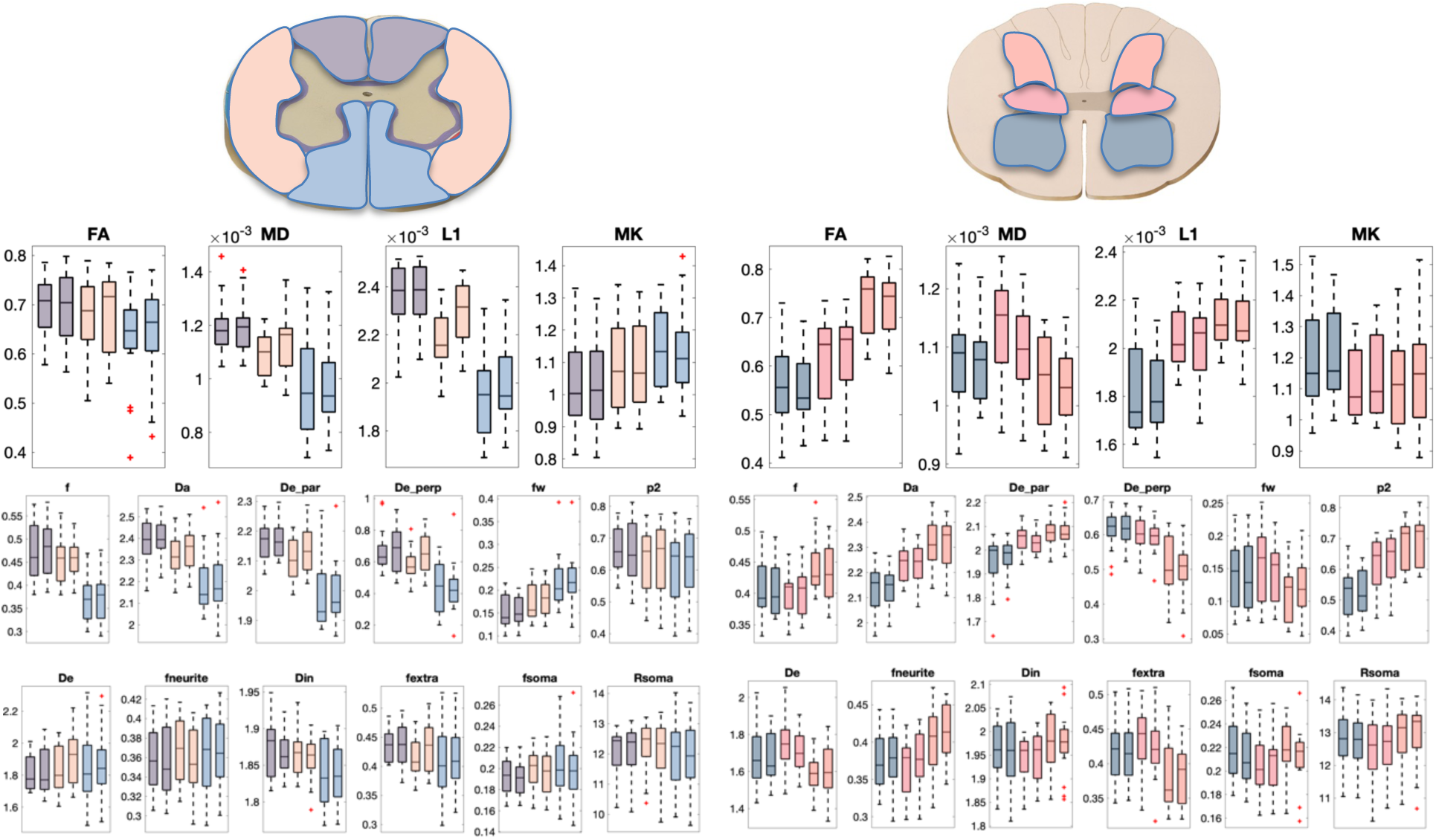
DTI (top), SMI (middle), and SANDI (bottom) show contrast within and between the somatotopically organized white and gray matter regions. Plots are colored based on location in the white matter (bilateral Dorsal Columns, Lateral Funiculi, Ventral Funiculi) and gray matter (dorsal horn, intermediate zone, and ventral horns).

## Discussion

We implemented a clinically feasible diffusion protocol for the in vivo human brain and spinal cord that enables modeling of the white and gray matter tissue microstructure of a large portion of the central nervous system. Specifically, this protocol enables DTI, the standard model of white matter (SMI), and its extension to gray matter via the SANDI model. We show that derived microstructural measures exhibit reasonable and expected tissue contrast in the brain and cord, with contrast across and within both white and gray matter of both structures. Notably, this study also represents the first images of SMI and SANDI modeling of the spinal cord, and the resulting contrasts and values are reasonable and within expectations given knowledge of brain data. However, the partial volume effects and small intra-cord structures provides insight into the current challenges and requirements for improvements to better characterize the entire central nervous system.

### Acquisition and considerations

The data acquisition takes approximately 16 minutes for each anatomy (brain and cord), and includes 6 b-values, 176 total diffusion weighted images, and 24 b=0 images. While we implement DTI, SMI, and SANDI, this acquisition also enables commonly employed multicompartment models that require only multiple shells using standard PGSE sequences, for example Neurite Orientation Dispersion and Density Imaging (NODDI) [19], Freewater DTI [54], Spherical Mean Technique (SMT) [21], White Matter Tract Imaging (WMTI) [55], among others.

The protocol was inspired by optimization performed by Schiavi et al. [49], who optimized a protocol for the human brain for the SANDI model, within the constraints of their 3T scanner equipped with 80mT/m gradients. By generating simulated tissue substrates, they assess bias and reproducibility of various acquisition schemes, ranging from 14 shells down to 6, and showing that at realistic noise levels (SNR = 100 after spherical averaging over all directions) reducing from 14 to 6 shells has no major impact on parameter estimation. However, a major challenge to acquisition on these systems is the long diffusion times, δ =24 ms and Δ =51 ms, which lead to an increased TE (and decreased SNR) but also an increased sensitivity to exchange, which may violate assumptions of the model and bias parameter estimation. Optimistically, with exchange between compartments on the scale of ∼20-50ms (as estimated in rat brain cortex [56]), biases due to exchange are lower than the impact of noise, however, biases still range on the order of 5-15% for most parameters. Similarly, the authors [49], and others [57, 58], highlight that measures such as Din are neither accurate nor precise without alternative acquisitions (linear and planar encodings, or varying diffusion times [59]), which matches our results where Din has lowest scan-rescan reproducibility in both the brain and cord.

While the protocol matched that of Schiavi et al. [49] in the brain, several innovations were needed in the spinal cord. Typical spinal cord diffusion protocols suggest cardiac triggering during the same point in the cardiac cycle to minimize susceptibility to motion effects [33], however, this reduces scan efficiency and would result in tremendously long scan times to acquire ∼200 total image volumes. Recent work [36] suggests that removing cardiac triggering, which enables acquiring ∼60% more data, combined with denoising, motion correction, and outlier removal and replacement, results in measures of similar DTI indices, with similar reproducibility as triggered acquisitions. In this work, we removed triggering (which also enables harmonization between brain and cord by keeping TR similar), applied MPPCA denoising [39], used spinal cord-specific motion correction [46] with slice-wise regularization along the cord and grouping of 8 successive low SNR volumes to improve robustness, and finally implemented a Patch2Self denoising algorithm that replaces outlier signals (i.e. slice dropouts) [43, 60, 61]. We note that we did not apply distortion correction, as these algorithms have not been fully optimized in the cord and do not quantitatively nor qualitatively improve derived metrics [62–64]. Overall, the acquisition and subsequent preprocessing resulted in high-fidelity images with white and gray matter contrast in diffusion weighted images, and facilitated DTI, SMI, and SANDI-based modeling on a clinical scanner.

### Brain and Cord

A major innovation in this work is the feasibility of a harmonized acquisition that will enable characterization and quantification of diffusion-based measures of soma and neurite densities and organizations, and possible biomarkers, throughout the neuroaxis from brain to cervical cord. Most studies of the central nervous system study the brain or cord in isolation, whereas both structures may contribute to clinical deficits. We hypothesize that the ability to simultaneously examine both will advance our understanding of various central nervous system pathologies.

One such example is multiple sclerosis (MS). Conventional MRI facilitates lesion visualization and quantification, in both brain and cord, yet lacks specificity to axonal content, both within lesions and in normal appearing tissue. Because of this, conventional T1 or T2 weighted measures of lesion load in the brain or in the cord only reveal low to moderate associations with cognitive or motor functions [65–67]. Utilizing advanced diffusion MRI to provide measures of neurite and soma organization, size, and density, and subsequent neurite/soma loss, swelling/shrinking, or edema may enable higher sensitivity to functional changes with the increased specificity to tissue microstructural changes.

Recent changes to the McDonald criteria for MS diagnosis (in 2017 and 2024) included dissemination of lesions in space by including both brain and spinal cord lesions (in structural images), with the goal of moving towards a biological diagnosis (as opposed to clinical means/symptoms); expanding the arsenal of tools to query tissue microstructure is a prerequisite to provide a more comprehensive picture of the disease.

Our results show some consistency between these two structures connected by the ascending and descending projection pathways. Using our protocol, the white matter of the cord exhibits anisotropies (FA, p2) within the range expected of white matter in the brain. Similar results are observed for diffusivities, both axonal and extra-axonal, as well as neurite fractions. Future work may investigate a pathway-specific analysis of the decussating and non-decussating corticospinal tracts to assess the feasibility of studying the length of this pathway from the cortex to the cord. Overall, our results show reasonable contrast from the brain to the cord, with values within expected ranges (despite differences in SNR and resolution) for white and gray matter microstructures.

### Multicompartment modeling of the spinal cord

This is the first application of the SMI and SANDI models in the in vivo human spinal cord. Previous works have characterized white and gray matter in the cord using NODDI and SMT, providing evidence of dispersion and neurite density changes in disease (MS) in both lesion and normal appearing tissue, as well as providing normative values metrics in the healthy cord[32, 68–71]. Here, we show that SMI and SANDI similarly provide contrast among the various white matter pathways and gray matter regions where heterogenous axonal environments are expected.

The advantage of SMI is that it encompasses a number of WM models made to capture Gaussian compartments, with axons represented by sticks [28]. While other models impose constraints on parameters, between parameters, or on forms of the fiber distribution to improve robustness, they may introduce biases into the parameter estimation. Here, supervised machine learning is used to improve precision of parameter estimates [28], while also providing microstructural maps without constraints or priors. We find neurite fractions of ∼.4-.5, axial diffusivity >2 (and greater than extra-axonal parallel diffusivity), orientation dispersion (p2) ∼0.7, and a relatively large partial volume fraction with free water (∼.15-.2). These maps are all in line with that observed in the brain white matter [28] (although with larger partial volume fraction with free water, and subsequently smaller neurite fraction), with maps that are anatomically and microstructurally feasible.

The advantage of the SANDI model is explicit modeling of compartments that are prevalent in the gray matter, in particular soma size and soma fraction [29]. This necessitates the high b-value data, for increased sensitivity to the small, restricted spherical compartments. Here, the disadvantage is the increased TE to reach this diffusion weighting, decreasing signal and increasing diffusion times. SMI, above, may benefit from a multi-shell acquisition that is limited to a b-value of ∼2000-3000 instead of occurring the signal loss associated with also modeling SANDI. The increased diffusion times increase biases due to exchange, for which there is no consensus on the time scale in the cord, and limits sensitivity to smaller soma. Despite this, this model also results in an intracellular diffusivity slightly greater than extracellular parallel diffusivity, and a neurite fraction of ∼.3-.45.

On the optimistic side, both models result in quantitative differences across both white and gray matter tissues, which is expected in the cord. For example, differences in locations of sensory synapses (dorsal horn) and motor cell bodies (ventral horn), along with branching nerve roots are expected to result in different microstructural environments across these structures. Moreover, reproducibility (particularly TRV) of most measures derived from SMI/SANDI was similar to that of DTI, with generally similar values between brain and cord. However, ICC was (in general) lower in the cord than brain. Some of this can be attributed smaller voxel volume in the cord, smaller tissues of interest and potentially incompletely mitigated motion.

Similarly, for white and gray matter contrast, partial volume effects from the small size and incompletely mitigated motion (across volumes), geometric distortion, and atlas-based tissue segmentation play a role. Together, this makes localization of specific changes in the small (often less than the size of a voxel) ascending/descending pathways or gray matter horns challenging, losing some of the spatial specificity but gaining mictostructural specificity gained with multi-compartment modeling. Another challenge is the large axons of the spinal cord, with a volume-weighted mean axon diameter often on the order of 2-5um depending on pathway, which may violate assumptions of a zero-radius cylinder (i.e., a stick) for the neurite compartment. Veraart et al. [72, 73], show that in vivo human brain, this assumption holds valid when using clinical scanners and subsequently long diffusion times, with the neurites indistinguishable from sticks. Similar studies should be performed in the cord to assess sensitivity to non-negligible axon radii.

### Limitations and Future

There are several limitations that must be acknowledged. First, is the limited sample size (N=11) of the current study. While 11 subjects (and N=5 rescans) is sufficient to assess reproducibility [49, 50, 74], it is not sufficient to provide comprehensive normative values across the population, nor is this a validation of the derived indices.

However, the goal was to demonstrate the potential of capturing newly-developed contrasts within and across tissue types on a clinical scanner. Second, these models may not be suitable for this acquisition – with these noise properties, diffusion times, and PGSE acquisition. In addition to biases described above due to long diffusion times, variance due to noise in parameter estimation, and possibly invalid stick-like assumptions in neurites, the use of alternative diffusion weightings (planar, spherical) may better condition model-fitting [28], and should be investigated in the future. Improvements in spinal cord image processing are also needed, including optimizing preprocessing associated with high b-value data, which challenges distortion and motion correction when most volumes are expected to be at the noise level. Finally, future studies should investigate the clinical utility of combining brain and cord features in studying neurological disorders, and use similar acquisitions to investigate the relationship between brain and cord.

## Conclusion

We have presented and evaluated a harmonized diffusion protocol to study the brain and spinal cord on a clinical scanner. The protocol enables DTI modeling, as well as advanced SMI and SANDI models to improve specificity in characterizing neurite and soma organization. We demonstrate feasibility of acquiring multi-shell diffusion data at high b-values and high SNR, show qualitatively reasonable contrast across brain and cord, and show SMI and SANDI have moderate reproducibility relative to DTI, with higher scan rescan variability in the cord than in the brain. Finally, we demonstrate that soma and neurite density with these models shows contrast across white matter pathways, and across gray matter regions in both structures, which is expected given known variation in neurite densities and diameters and cellular architectures and densities. This protocol can be employed in a reasonable amount of time to study CNS pathologies and investigate biomarkers of structural integrity throughout the neuroaxis.

## Acknowledgements

This work was supported by the National Institutes of Health (NIH) under award numbers NIH K01EB030039 (KO), 1R03TR004434 (KO), K01EB032898 (KS), 5R01NS109114 (SS), 5R01NS117816 (SS) and 5R01NS104149 (SS), R01EB017230 (BL).

## Author Contributions

KGS (writing, analysis, conceptualization); MP (analysis, experimental design); AAW (experimental design, data curation); KPOG (experimental design, data curation); MP (experimental design, analysis); BAL (funding, conceptualization, supervision); SAS (funding, conceptualization, supervision)

## Declaration of competing interest

The authors have no competing interests to declare.

## Ethics

All participants from whom data were used in this manuscript, provided written informed consent (and consent to publish) according to the declaration of Helsinki.

## Data and Code Availability

Data will be made available in BIDS format in openneuro sharing platform upon acceptance of the manuscript. To be updated with DOI.

## Notes

### Competing Interest Statement

The authors have declared no competing interest.

## References

1. Nieuwenhuys, R., J. Voogd, and C.v. Huijzen, The human central nervous system. 4th ed. 2008, New York: Springer. xiv, 967 p.

2. Felten, D., Netter’s atlas of neuroscience. 4. ed. 2021, Philadelphia: Elsevier, Inc. pages cm.

3. Le Bihan, D. and M. Iima, Diffusion Magnetic Resonance Imaging: What Water Tells Us about Biological Tissues. PLOS Biology, 2015. 13(7): p. e1002203.

4. Beaulieu, C., The basis of anisotropic water diffusion in the nervous system - a technical review. NMR Biomed, 2002. 15(7-8): p. 435–55.

5. Novikov, D.S., et al., Quantifying brain microstructure with diffusion MRI: Theory and parameter estimation. NMR Biomed, 2018: p. e3998.

6. Novikov, D.S., V.G. Kiselev, and S.N. Jespersen, On modeling. Magn Reson Med, 2018. 79(6): p. 3172–3193.

7. Basser, P.J., J. Mattiello, and D. LeBihan, MR diffusion tensor spectroscopy and imaging. Biophys J, 1994. 66(1): p. 259–67.

8. Pierpaoli, C., et al., Diffusion tensor MR imaging of the human brain. Radiology, 1996. 201(3): p. 637–48.

9. Jones, D.K. and M. Cercignani, Twenty-five pitfalls in the analysis of diffusion MRI data. NMR Biomed, 2010. 23(7): p. 803–20.

10. Jones, D.K., M.A. Horsfield, and A. Simmons, Optimal strategies for measuring diffusion in anisotropic systems by magnetic resonance imaging. Magn Reson Med, 1999. 42(3): p. 515–25.

11. Mori, S. and J. Zhang, Principles of diffusion tensor imaging and its applications to basic neuroscience research. Neuron, 2006. 51(5): p. 527–39.

12. Vedantam, A., et al., Diffusion tensor imaging of the spinal cord: insights from animal and human studies. Neurosurgery, 2014. 74(1): p. 1–8; discussion 8; quiz 8.

13. Kroenke, C.D., J.J. Ackerman, and D.A. Yablonskiy, On the nature of the NAA diffusion attenuated MR signal in the central nervous system. Magn Reson Med, 2004. 52(5): p. 1052–9.

14. Jespersen, S.N., et al., Modeling dendrite density from magnetic resonance diffusion measurements. Neuroimage, 2007. 34(4): p. 1473–86.

15. Sotiropoulos, S.N., T.E. Behrens, and S. Jbabdi, Ball and rackets: Inferring fiber fanning from diffusion-weighted MRI. Neuroimage, 2012. 60(2): p. 1412–25.

16. Reisert, M., et al., MesoFT: unifying diffusion modelling and fiber tracking. Med Image Comput Comput Assist Interv, 2014. 17(Pt 3): p. 201–8.

17. Jelescu, I.O., et al., Degeneracy in model parameter estimation for multi-compartmental diffusion in neuronal tissue. NMR Biomed, 2016. 29(1): p. 33–47.

18. Novikov, D.S., et al., Rotationally-invariant mapping of scalar and orientational metrics of neuronal microstructure with diffusion MRI. Neuroimage, 2018. 174: p. 518–538.

19. Zhang, H., et al., NODDI: practical in vivo neurite orientation dispersion and density imaging of the human brain. Neuroimage, 2012. 61(4): p. 1000–16.

20. Kaden, E., et al., Multi-compartment microscopic diffusion imaging. Neuroimage, 2016. 139: p. 346–359.

21. Kaden, E., F. Kruggel, and D.C. Alexander, Quantitative mapping of the per-axon diffusion coefficients in brain white matter. Magn Reson Med, 2016. 75(4): p. 1752–63.

22. Fieremans, E., et al., Novel white matter tract integrity metrics sensitive to Alzheimer disease progression. AJNR Am J Neuroradiol, 2013. 34(11): p. 2105–12.

23. Fieremans, E., J.H. Jensen, and J.A. Helpern, White matter characterization with diffusional kurtosis imaging. Neuroimage, 2011. 58(1): p. 177–88.

24. Coronado-Leija, R., et al., Volume electron microscopy in injured rat brain validates white matter microstructure metrics from diffusion MRI. ArXiv, 2024.

25. Fieremans, E., et al., Diffusion distinguishes between axonal loss and demyelination in brain white matter. Proc. Int. Soc. Magn. Reson. Med., 2012. 20.

26. Jelescu, I.O., et al., In vivo quantification of demyelination and recovery using compartment-specific diffusion MRI metrics validated by electron microscopy. Neuroimage, 2016. 132: p. 104–114.

27. Lee, H.H., et al., A time-dependent diffusion MRI signature of axon caliber variations and beading. Commun Biol, 2020. 3(1): p. 354.

28. Coelho, S., et al., Reproducibility of the Standard Model of diffusion in white matter on clinical MRI systems. Neuroimage, 2022. 257: p. 119290.

29. Palombo, M., et al., SANDI: A compartment-based model for non-invasive apparent soma and neurite imaging by diffusion MRI. Neuroimage, 2020. 215: p. 116835.

30. Genc, S., et al. Repeatability of Soma and Neurite Metrics in Cortical and Subcortical Grey Matter. in Computational Diffusion MRI. 2021. Cham: Springer International Publishing.

31. Schiavi, S., et al., Mapping tissue microstructure across the human brain on a clinical scanner with soma and neurite density image metrics. Hum Brain Mapp, 2023. 44(13): p. 4792–4811.

32. Schilling, K.G., et al. Investigating multi-compartment diffusion MRI models in the cervical spinal cord of multiple sclerosis patients. in International Society of Magnetic Resonance in Medicine. 2021.

33. Cohen-Adad, J., et al., Generic acquisition protocol for quantitative MRI of the spinal cord. Nat Protoc, 2021.

34. Held, P., et al., MRI of the abnormal cervical spinal cord using 2D spoiled gradient echo multiecho sequence (MEDIC) with magnetization transfer saturation pulse. A T2* weighted feasibility study. J Neuroradiol, 2003. 30(2): p. 83–90.

35. Wilm, B.J., et al., Reduced field-of-view MRI using outer volume suppression for spinal cord diffusion imaging. Magn Reson Med, 2007. 57(3): p. 625–30.

36. Kurt, G.S., et al., Influence of preprocessing, distortion correction and cardiac triggering on the quality of diffusion MR images of spinal cord. bioRxiv, 2023: p. 2023.09.26.559530.

37. Fischl, B., FreeSurfer. Neuroimage, 2012. 62(2): p. 774–81.

38. Fischl, B., et al., Whole brain segmentation: automated labeling of neuroanatomical structures in the human brain. Neuron, 2002. 33(3): p. 341–55.

39. Veraart, J., et al., Denoising of diffusion MRI using random matrix theory. Neuroimage, 2016. 142: p. 394–406.

40. Kellner, E., et al., Gibbs-ringing artifact removal based on local subvoxel-shifts. Magn Reson Med, 2016. 76(5): p. 1574–1581.

41. Andersson, J.L., S. Skare, and J. Ashburner, How to correct susceptibility distortions in spin-echo echo-planar images: application to diffusion tensor imaging. Neuroimage, 2003. 20(2): p. 870–88.

42. Andersson, J.L.R., et al., Incorporating outlier detection and replacement into a non-parametric framework for movement and distortion correction of diffusion MR images. Neuroimage, 2016. 141: p. 556–572.

43. Fadnavis, S., J. Batson, and E. Garyfallidis, Patch2Self: denoising diffusion MRI with self-supervised learning. arXiv preprint arXiv:2011.01355, 2020.

44. De Leener, B., et al., SCT: Spinal Cord Toolbox, an open-source software for processing spinal cord MRI data. Neuroimage, 2017. 145(Pt A): p. 24–43.

45. Veraart, J., E. Fieremans, and D.S. Novikov, Diffusion MRI noise mapping using random matrix theory. Magn Reson Med, 2016. 76(5): p. 1582–1593.

46. Xu, J., et al., Improved in vivo diffusion tensor imaging of human cervical spinal cord. Neuroimage, 2013. 67: p. 64–76.

47. Jensen, J.H., et al., Diffusional kurtosis imaging: the quantification of non-gaussian water diffusion by means of magnetic resonance imaging. Magn Reson Med, 2005. 53(6): p. 1432–40.

48. Reisert, M., et al., Disentangling micro from mesostructure by diffusion MRI: A Bayesian approach. Neuroimage, 2017. 147: p. 964–975.

49. Schiavi, S., et al., Dissecting brain grey and white matter microstructure: a novel clinical diffusion MRI protocol. bioRxiv, 2022: p. 2022.04.08.487640.

50. Veraart, J., et al., The variability of MR axon radii estimates in the human white matter. Hum Brain Mapp, 2021. 42(7): p. 2201–2213.

51. Palombo, M., et al., Joint estimation of relaxation and diffusion tissue parameters for prostate cancer with relaxation-VERDICT MRI. Sci Rep, 2023. 13(1): p. 2999.

52. Schober, P., C. Boer, and L.A. Schwarte, Correlation Coefficients: Appropriate Use and Interpretation. Anesth Analg, 2018. 126(5): p. 1763–1768.

53. Salarian, A., Intraclass Correlation Coefficient (ICC). 2024.

54. Pasternak, O., et al., Free water elimination and mapping from diffusion MRI. Magn Reson Med, 2009. 62(3): p. 717–30.

55. Jelescu, I.O., et al., One diffusion acquisition and different white matter models: how does microstructure change in human early development based on WMTI and NODDI? Neuroimage, 2015. 107: p. 242–256.

56. Jelescu, I.O., et al., Neurite Exchange Imaging (NEXI): A minimal model of diffusion in gray matter with inter-compartment water exchange. Neuroimage, 2022. 256: p. 119277.

57. Dhital, B., et al., Intra-axonal diffusivity in brain white matter. Neuroimage, 2019. 189: p. 543–550.

58. Howard, A.F., et al., Estimating axial diffusivity in the NODDI model. Neuroimage, 2022. 262: p. 119535.

59. Coelho, S., et al., Resolving degeneracy in diffusion MRI biophysical model parameter estimation using double diffusion encoding. Magn Reson Med, 2019. 82(1): p. 395–410.

60. Schilling, K.G., et al., Patch2Self denoising of diffusion MRI in the cervical spinal cord improves intra-cord contrast, signal modelling, repeatability, and feature conspicuity. medRxiv, 2021: p. 2021.10.04.21264389.

61. Pizzolato, M., et al., Axial and radial axonal diffusivities and radii from single encoding strongly diffusion-weighted MRI. Med Image Anal, 2023. 86: p. 102767.

62. Snoussi, H., et al. Geometric Evaluation of Distortion Correction Methods in Diffusion MRI of the Spinal Cord. in 2019 IEEE 16th International Symposium on Biomedical Imaging (ISBI 2019). 2019.

63. Snoussi, H., et al., Evaluation of distortion correction methods in diffusion MRI of the spinal cord. arXiv preprint arXiv:2108.03817, 2021.

64. Schilling, K.G., et al., Influence of preprocessing, distortion correction and cardiac triggering on the quality of diffusion MR images of spinal cord. Magn Reson Imaging, 2024. 108: p. 11–21.

65. Hackmack, K., et al., Can we overcome the’clinico-radiological paradox’ in multiple sclerosis? J Neurol, 2012. 259(10): p. 2151–60.

66. Mollison, D., et al., The clinico-radiological paradox of cognitive function and MRI burden of white matter lesions in people with multiple sclerosis: A systematic review and meta-analysis. PLoS One, 2017. 12(5): p. e0177727.

67. Johnen, A., et al., Resolving the cognitive clinico-radiological paradox - Microstructural degeneration of fronto-striatal-thalamic loops in early active multiple sclerosis. Cortex, 2019. 121: p. 239–252.

68. Grussu, F., et al., Neurite orientation dispersion and density imaging of the healthy cervical spinal cord in vivo. Neuroimage, 2015. 111: p. 590–601.

69. By, S., et al., Application and evaluation of NODDI in the cervical spinal cord of multiple sclerosis patients. Neuroimage Clin, 2017. 15: p. 333–342.

70. Grussu, F., et al., Neurite dispersion: a new marker of multiple sclerosis spinal cord pathology? Ann Clin Transl Neurol, 2017. 4(9): p. 663–679.

71. By, S., et al., Multi-compartmental diffusion characterization of the human cervical spinal cord in vivo using the spherical mean technique. NMR Biomed, 2018. 31(4): p. e3894.

72. Veraart, J., E. Fieremans, and D.S. Novikov, On the scaling behavior of water diffusion in human brain white matter. Neuroimage, 2019. 185: p. 379–387.

73. Veraart, J., et al., Nonivasive quantification of axon radii using diffusion MRI. Elife, 2020. 9.

74. Koller, K., et al., MICRA: Microstructural image compilation with repeated acquisitions. Neuroimage, 2021. 225: p. 117406.

